# Neuronal activity and learning in local cortical networks are modulated by the action-perception state

**DOI:** 10.1101/537613

**Authors:** Ben Engelhard, Ran Darshan, Nofar Ozeri-Engelhard, Zvi Israel, Uri Werner-Reiss, David Hansel, Hagai Bergman, Eilon Vaadia

**Affiliations:** Department of Medical Neurobiology, Institute of Medical Research Israel-Canada, The Hebrew University-Hadassah Medical School, Jerusalem, Israel; Edmond and Lily Safra Center for Brain Sciences, The Interdisciplinary Center for Neural Computation, The Hebrew University, Jerusalem, Israel; Laboratoire de Neurophysique et Physiologie UMR8119-CNRS, Université René Descartes, Paris, France; Center for Function and Restorative Neurosurgery, Department of Neurosurgery, Hadassah University Hospital, Jerusalem, Israel; The Alexander Silberman Institute of Life Sciences, The Hebrew University, Jerusalem, Israel

## Abstract

During sensorimotor learning, neuronal networks change to optimize the associations between action and perception. In this study, we examine how the brain harnesses neuronal patterns that correspond to the current action-perception state during learning. To this end, we recorded activity from motor cortex while monkeys either performed a familiar motor task (movement-state) or learned to control the firing rate of a target neuron using a brain-machine interface (BMI-state). Before learning, monkeys were placed in an observation-state, where no action was required. We found that neuronal patterns during the BMI-state were markedly different from the movement-state patterns. BMI-state patterns were initially similar to those in the observation-state and evolved to produce an increase in the firing rate of the target neuron. The overall activity of the non-target neurons remained similar after learning, suggesting that excitatory-inhibitory balance was maintained. Indeed, a novel neural-level reinforcement-learning network model operating in a chaotic regime of balanced excitation and inhibition predicts our results in detail. We conclude that during BMI learning, the brain can adapt patterns corresponding to the current action-perception state to gain rewards. Moreover, our results show that we can predict activity changes that occur during learning based on the pre-learning activity. This new finding may serve as a key step toward clinical brain-machine interface applications to modify impaired brain activity.

## Introduction

Brain-machine interfaces (BMIs) are powerful tools initially developed to read brain activity and translate it into actions [1–7] in order to restore motor function in paralyzed patients [8,9]. These studies also facilitated understanding of sensorimotor control and learning by analyzing the relationships between neural activity and the resulting action. Recently, another class of BMI algorithms, based on modulation of neuronal activity by operant conditioning, has gained interest. This idea was first demonstrated almost 50 years ago, when monkeys were trained to change the firing rate of a single neuron [10]. As simultaneous recording of many neurons became possible, operant neural conditioning emerged as a powerful approach to modulating the *neuronal state* (as expressed by firing rates, oscillations, and correlations) and examining how it changes the *action-perception state* of the subject (AP state) [11–18]. We define an AP state as a particular relationship between actions and their perceptual consequences [19]. This approach can be applied translationally to detect and modulate abnormal brain activity, thereby using neural conditioning to ameliorate behavioral disorders [20–22].

These findings raised fundamental questions about learning: (1) How is the neural state linked to the learning process and the resulting modified AP state? (2) How limited are the possible neural states in a given local circuit? For example, is the pattern of neuronal co-modulations fixed for a given network, or does it change in different AP states?

To approach these questions, we combined experimental and modeling tools and explored state-dependent learning in a local circuit of the motor cortex in monkeys. Experimentally, we used the BMI setting to condition the firing rate of a single (target) neuron, making it the only causal variable leading to reward. In parallel, we simultaneously recorded many non-target neurons in the local circuit. Our modeling work was aimed at finding the theoretical underpinnings of learning in the local circuit and the functional mechanisms by which it is achieved.

Our model addresses two essential components of reinforcement learning (RL): variability in network activity, required for exploration, and plasticity, required for learning. RL models provide a high-level description of how behavioral learning is shaped by rewards [23,24]. However, the neural implementation of such learning remains an open question. Cortical variability has been proposed to emerge in a regime of balanced excitation and inhibition (EI), where the network can exhibit chaotic dynamics [25–27]. Here, we develop a novel model implementing reward-mediated plasticity in a chaotic EI-balanced network with slow synapses [28]. We show how plasticity that drives learning can lead to large network reorganizations while maintaining the ability to drive exploration, thus providing a mechanistic implementation of reinforcement-based learning in a cortical network.

As networks learn to volitionally control the activity of a single neuron, we show that activity in both the biological network and the model evolve from the pre-learning neural state to a new neural state, which enables increased firing of the target neuron. In addition, these pre-learning states permit prediction of the upcoming network reorganization during learning.

## Results

### Measuring neuronal activity in different action-perception states

Two monkeys were implanted with 96-electrode arrays in the arm area of the primary motor cortex. We probed network activity across action-perception states in experimental sessions consisting of four blocks (Figure 1A). The first block, the *movement-state*, consisted of a familiar center-out task, where monkeys performed hand grip and reaching movements to one of eight directions chosen randomly on each trial (Figure 1C, left). In the second block (*observation-state*), the monkeys’ arms were comfortably restrained, and a cursor moved in each trial from the center of the screen (origin) toward one of the eight targets used in the movement-state block. However, the distance of the cursor from the origin to the target was chosen randomly every 100 ms, so rewards were delivered randomly as well (when the cursor randomly reached the target; Figure 1C, middle). The third block was the *BMI-state*. Here, conditions were similar to the observation-state, but one target (chosen at random per session) was fixed for all trials of the block, and the position of the cursor at time *t* depended on the firing rate of a single neuron (target neuron) in a window of 400 ms preceding *t* (Figure 1C, right); when the firing rate was high enough, the cursor reached the target, and a liquid reward was delivered. In most sessions (24/34), the monkeys learned to reliably increase the firing rates of single neurons in the BMI-state (examples in Figure 1B). The remaining sessions were not analyzed further. By design, the overt behavioral context for the monkeys (both hands restrained, cursor moving one-dimensionally on the screen) was similar between the observation- and BMI-states, thus separating the BMI-state context from the movement-state context via an intermediate state. The last block of the session was a repeat of the movement-state after BMI learning.

**Figure 1.**
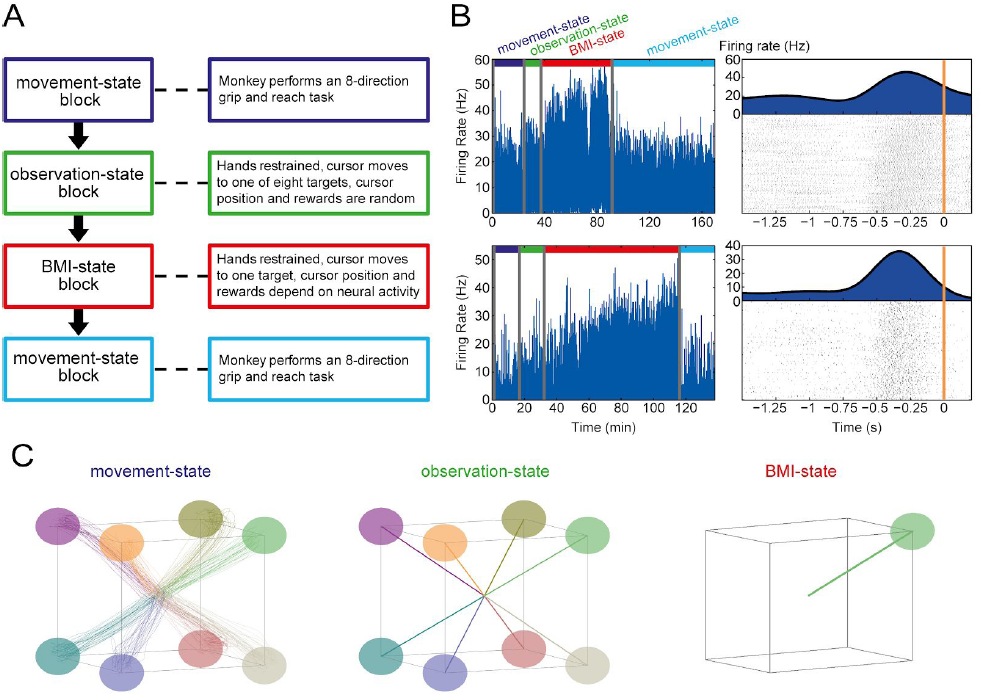
Behavioral task. (A) Experimental block design of a single session. (B) Left panels show examples of the firing rate of the target neuron across two complete sessions for both monkeys; the firing rate increased during the BMI-state block. Right panels show raster plots for the same neurons during 400 successful trials in the BMI-state block. Rasters are locked to reward (time 0, orange line). Above the rasters, the Peri-Event Time Histogram (PETH) depicts the increase in firing rate that led to reward. (C) Cursor trajectories during all trials of one session (in each trial the there was a single target) Left: trajectories during the movement-state block. Center: trajectories during the observation-state block. Right: trajectories during the BMI-state block.

### Population activity depends on the action-perception state

To investigate possible changes in population activity during different action-perception states, we first used state-space analysis, which permits visualization of high-dimensional neural data. We performed Gaussian-process factor analysis (GPFA) [29] on the *non-target* neurons during trials of all four blocks in the session (see Methods). Representative examples of the population activity in neural state-space from three single sessions (Figure 2A) demonstrate three features of state-dependent population activity. (1) Activity during different action-perception states occupies distinct subsets of neural space. (2) Activity in the two movement-state blocks (first and last blocks, blue) is overlaid in neural space, indicating that neural drift throughout the session is an unlikely cause of this separation. (3) Activity during the BMI-state (red) appears closer to that of the observation-state (green) than to that of either of the movement-state blocks.

**Figure 2.**
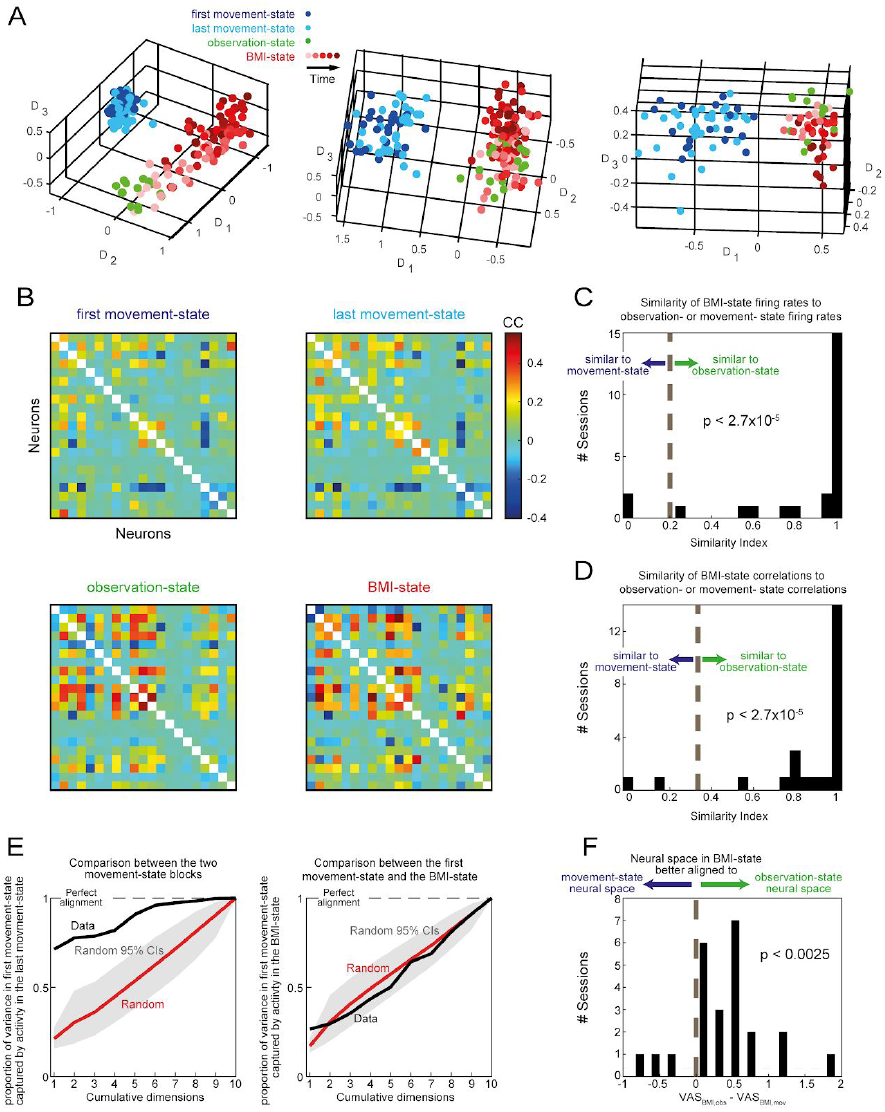
Comparison of population activity in the different action-perception states. (A) The panel shows the neural state-space during the different behavioral blocks in three complete sessions. Each point represents a trial. For clarity, every 5th trial is shown in the plots. Trials are color coded according to the block in which they occurred: dark and light blue: first and last movement-state blocks. Green: observation-state block. Light to dark red: BMI-state block. For this block, the color gradient also shows the temporal order of the trials along the session. The axes represent the first three dimensions of the neural state, computed using GPFA (see Methods). (B) Correlation matrices between all pairs of neurons recorded in one session, for the different behavioral blocks. Each bin represent the correlation between two neurons, and each matrix shows the structure of the pairwise correlations of the same set of neurons in one block. Note the difference between correlations during the movement-states and BMI-state. (C) Similarity index to the average firing rate vectors during BMI-state in all sessions (see Methods). High values indicate increased similarity to the observation-state firing rates, and lower values increased similarity to the movement-states firing rates. Chance level (equal similarity) is indicated by the dashed line. Significance is denoted in the figure (based on a Wilcoxon signed rank test of the median of the distribution to the chance level). (D) Similar to C, but showing the similarity index of the neural correlation matrices. (E) Variance alignment analysis for same session shown in B. The neural spaces in the two movement-state blocks are significantly aligned (left panel), while alignment between the first movement-state and the BMI-state is not different than random (right panel). (F) Similarity to the BMI-state neural space computed by variance alignment analysis (all sessions). For each session we subtracted the alignment score between the movement- and BMI-states from the alignment score of the observation- and BMI-states. A positive value thus indicates a higher alignment score between the observation- and BMI-states. The histogram is significantly skewed to positive values (p<0.0025, Wilcoxon signed rank test).

We quantified these three features of the neuronal population activity using the *full dimensionality* of the data in all sessions. To quantify observations 1 and 2, we used a statistical test on the firing rate distributions in the different blocks, based on the Euclidean distances between and within the distributions and the Kolmogorov-Smirnov statistic (see Methods). We found that in most sessions (21/24), firing rates in the two movement-state blocks were not significantly different, whereas in all sessions, firing rates in the BMI-state were significantly different from those in the movement-state blocks. Additionally, in most sessions (23/24), firing rates in the observation-state blocks were significantly different from those in the movement-state blocks (p<0.01, after Bonferroni correction for the number of sessions).

Next, we tested if similar state-dependent features are exhibited in the co-modulation pattern of the neurons. The correlation matrices for all non-target neurons in one recording session changed dynamically depending on the action-perception state (Figure 2B). The similar correlation patterns in the first and last movement-state blocks are dramatically different from the correlation patterns in the observation- and BMI-state blocks, which are quite similar to each other. To quantify this observation for all sessions, we used a statistical test comparing the difference between correlation matrices in different blocks to the difference in correlations within each block where two halves of data (sampled randomly) were compared to each other (see Methods). We found that correlation matrices in the two movement-state blocks were not significantly different from each other in any session, whereas in almost all sessions (23/24), the correlations during the BMI-state were significantly different from the correlations in either movement-state block (p<0.01, after Bonferroni correction for the number of sessions). We conclude that correlations can change significantly depending on the action-perception state, and these changes do not appear to be the result of neural or measurement drift, given that the two movement-state blocks are furthest from each other in time.

The similarity between the correlation patterns during the observation- and BMI-states supports the notion expressed above that activity during the BMI-state is closer to that of the observation-state than to that of either movement-state block. We quantified this feature by examining the similarities in firing rates and correlations matrices between states. To do so, we calculated a similarity index for each session based on the euclidean distance between activity in the BMI-state and activity in the other states (using either firing rates or correlations as the relevant measure; see Methods). The results (Figure 2C,D) show that for both measures, across all sessions, activity during the BMI-state was significantly more similar to the activity during the observation-state than that of either of the movement-state blocks (p<2.7×10^-5^ for firing rates and p<2.7×10^-5^ for correlations, Wilcoxon signed rank test). These analyses took into account the possibility that the monkeys were using a re-aiming strategy during the BMI-state block (repurposing the movement-state patterns; see Methods). We conclude that during BMI learning, the network initiates exploration in dimensions of patterns of covariance in the most recent state (the observation-state in our experiment), rather than exploring for patterns from the repertoire of previously learned movement-space.

To directly test the relationships among the neural spaces explored by the network during the different blocks, we performed variance alignment analysis [30], which computes an alignment score between two neural spaces that is agnostic to scaling or rotations. This analysis, conducted for the same session as Figure 2B, shows that the two movement-state blocks share the same neural space, which is different from the BMI-state neural space (Figure 2E). To test further if the neural space during the BMI-state was more similar to the observation-state than that of the movement-state, we computed for each session the variance alignment score between the BMI- and observation-states (VAS_BMI,obs_) and the variance alignment score between the BMI and the first movement-state (VAS_BMI,mov_). We then plotted the distribution of the differences in these scores across sessions: VAS_BMI,obs_-VAS_BMI,mov_. Positive numbers indicate higher similarity between the BMI- and observation-states spaces, while negative numbers indicate higher similarity between the BMI- and the first movement-state spaces. The histogram is highly skewed to positive values (Figure 2F), indicating a significantly higher similarity between the observation- and BMI-state neural spaces (p<0.0025, Wilcoxon signed rank test).

Thus, overall, these analyses showed that the correlation structure and low-dimensional representation can change significantly as a function of the action-perception state. In addition, neuronal activity (as reflected by firing rates, pairwise correlations, and the neural state space) during BMI learning is different from that occurring during movements. Finally, activity during BMI learning is more similar to that occurring during the observation-state than to that occurring during movements. These results further support the notion that activity during BMI learning may have evolved from the pre-learning (observation-state) neural patterns.

### Learning in single neurons depends on the pre-learning neural correlation pattern

To challenge this interpretation, we investigated the dynamics in the local network by studying changes in neural activity between the observation-state and the BMI-state in more detail. To test the effects of learning on the population, we first assessed the significance of the changes in firing rates for all non-target neurons between the observation- and BMI-states (Wilcoxon rank-sum test, p<0.05). For each neuron, we quantified a normalized measure termed the Change in Firing Rate Index (ΔFR_indx_) as follows:

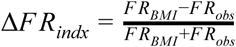

where *FR*_*BMI*_, *FR*_*obs*_ are the average firing rates in the 800 ms preceding reward in the BMI- and observation-state blocks, respectively.

The distribution of ΔFR_indx_ across the population (Figure 3A) shows that 64% (184/287) of the non-target neurons changed their firing rate significantly during BMI learning, suggesting that learning-related changes occurred at a large scale. These changes were not due to noise or neuronal drift, as they were significantly larger than changes between the two movement-state blocks (p<2.3×10^-5^, Brown–Forsythe test, Figure S1). Additional analyses (Figure S2) suggested that the activity patterns elicited during BMI learning were not related to either the sensory stimulus or physical movements.

**Figure 3.**
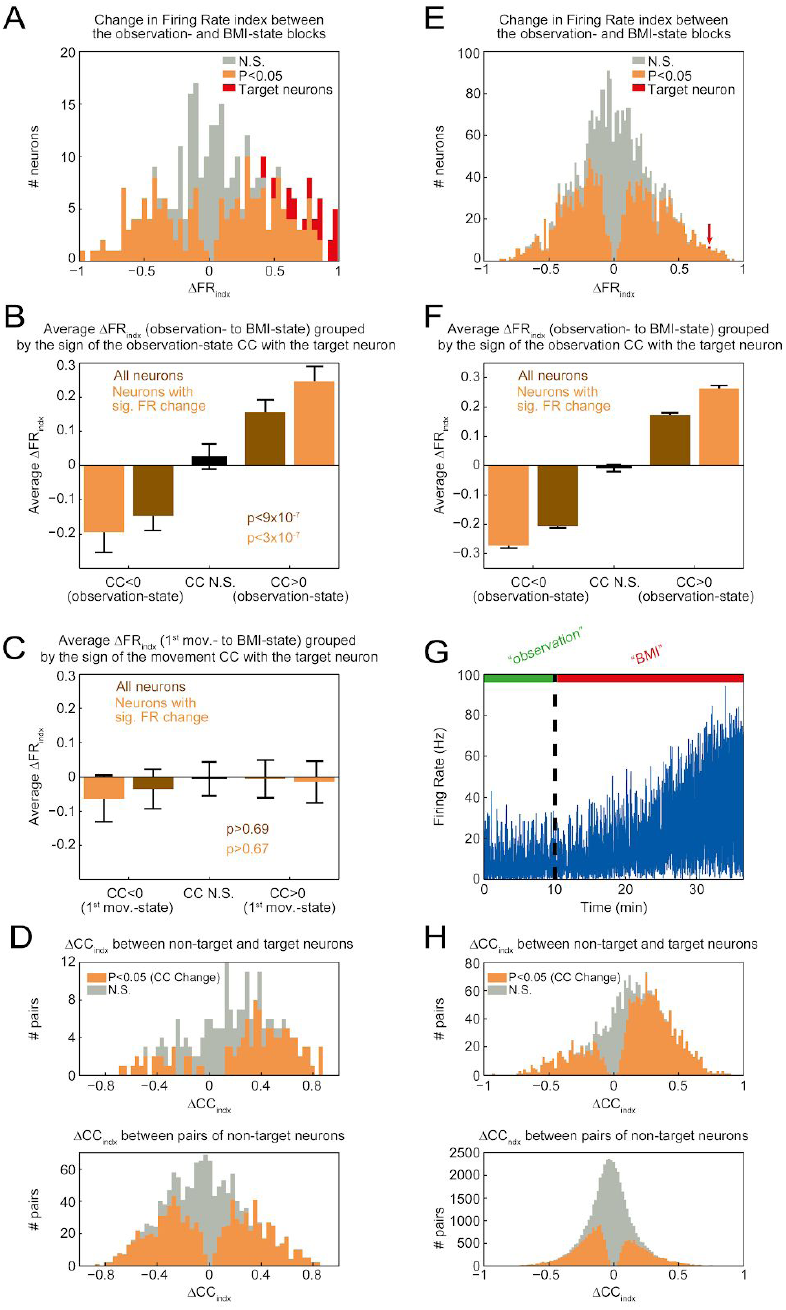
Activity changes across the network during learning, experiment and model. (A) Change in Firing Rate index (ΔFR_indx_) between the observation and BMI-state blocks (all sessions and neurons). Target neurons are marked in red, non-target neurons in gray (non-significant change in firing rate) or orange (significant change in firing rate, p<0.05, Wilcoxon rank-sum test). (B) Average ΔFR_indx_ grouped by the sign of the CC with the target neuron during the observation-state block. For all neurons (brown), neurons which evidenced a significant change in firing rate (orange) and neurons with non-significant CC with the target neuron (black). Error bars are s.e.m. (C) Same as B, but this time ΔFR_indx_ is calculated between the first movement-state block (1st mov.-state) and the BMI-state block, and the grouping is by the sign of the CC with the target neuron during the first movement-state block. (D) ΔCC_indx_ between neuronal pairs. Top: between non-target and target neurons. Bottom: between pairs of non-target neurons. Orange color denotes significant change in the CC between the two blocks (p<0.05, two-tailed z-test on the Fisher transformed CCs). Gray color denotes no significant change in the CCs. (E) Change in Firing Rate index (ΔFR_indx_) between the “observation” (pre-learning) and “BMI” states (learning) blocks in the simulation. The target neuron is marked in red and indicated by the red arrow, non-target neurons in gray (non-significant change in firing rate) or orange (significant change in firing rate, p<0.05, Wilcoxon rank-sum test). (F) Average ΔFR_indx_ grouped by the sign of the CC with the target neuron during the “observation” state block in the simulation. For all neurons (brown), neurons which evidenced a significant change in firing rate (orange) and neurons with non-significant CC with the target neuron (black). Error bars are s.e.m. (G) Firing rate of the target neuron during the simulated session. Learning started at t=10 min. (beginning of the “BMI” block). (H) ΔCC_indx_ between neuronal pairs in the simulation. Top: between non-target and target neurons. Bottom: between pairs of non-target neurons. Orange color denotes significant change in the CC between the two blocks (p<0.05, two-tailed z-test on the Fisher transformed CCs). Gray color denotes no significant change in the CCs.

Interestingly, although our learning task called for a rate *increase* of the target neuron, we did not observe an overall increase of firing rates across the population; the fraction of neurons that significantly increased or decreased their firing rate was similar (56% 103/184 sig. increased, 44%, 81/184 sig. decreased) and not significantly different from an equal division (p>0.12, binomial test). Thus, learning affected most neurons in a balanced manner across the population.

Can we predict activity changes following learning at the single-neuron level? If indeed the brain harnesses the observation-state patterns to effectuate learning, these patterns should be related to the observed changes in activity. Because successful task performance explicitly required increasing the firing rate of the target neuron, we looked at the correlation coefficients (CCs) of the firing rates between the target neuron and all non-target neurons during the observation-state block, and related these correlations to the firing rate changes of the neurons due to learning (see Methods). We found that most of the non-target neurons (61%, 176/287) had CCs with the target neuron that were significantly different from zero (p<0.01, two-tailed t-test) during the observation-state. The sign of these CCs was clearly related to the firing rate changes (ΔFR_indx_; Figure 3B). Non-target neurons that had negative CCs with the target neuron during the observation-state tended to decrease their firing rate due to learning, whereas those neurons that were positively correlated with the target neuron during the observation-state tended to increase their firing rate. The difference in firing rate changes between these two groups was highly significant (p<9×10^-7^, Wilcoxon rank-sum test). This result shows that learning-related changes across the network were strongly related to a specific feature of the observation-state correlations pattern: the correlations between the non-target neurons and the neuron that drives behavior during BMI. On average, neurons with a non-significant pre-learning CC with the target neuron did not change their firing rates (Figure 3B). Thus, by calculating the pre-learning neural correlation patterns related to the upcoming task-relevant (target) neuron, we could predict at 72% accuracy whether a neuron in the local population that was not causally related to the task would increase or decrease its firing rate. In contrast, no such relationship existed between the learning-related changes in the neural correlation pattern during the first (Figure 3C) or last (Figure S1) movement-state blocks. This result highlights the state-dependent nature of the learning process and supports the idea that the network adapts its activity patterns from the current state at the start of learning in order to accomplish the task.

We also attempted to predict the neural correlation pattern during BMI learning based on the pre-learning pattern. To study this question, we compared the CCs between pairs of non-target neurons (n=1947) in the different AP states. The correlation between the CCs during the observation- and BMI-states was 0.69. In contrast, the correlation between CCs during the BMI-state and the first movement-state block was 0.2, supporting once again the idea that activity patterns during BMI learning evolved from the observation-state patterns.

The similarity between the observation- and BMI-states could imply that there was little or no change in the CCs between these two states. Alternatively, widespread changes in the individual CCs could occur despite similarity in the overall *pattern* of correlations (e.g. CCs mostly maintain their sign and their relative strength). To evaluate these alternatives, we calculated the number of CCs that significantly changed between the observation- and BMI-states. We found that 74% of pairs significantly changed their CCs (n=1443/1947, p<0.05, two-tailed z-test on the Fisher-transformed CCs), but for most pairs CCs maintained their sign (72%, n=1413/1947). Thus, we find widespread changes in individual CCs with maintenance of the overall correlation pattern. To investigate how these changes in CCs are distributed across the population, we defined a measure to determine the changes in amplitude of the CCs as follows:

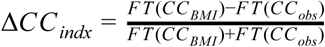

where *CC*_*BMI*_, *CC*_*obs*_ are the correlation coefficients for a given neuronal pair in the BMI- and observation-states, respectively, and *FT* (*x*) denotes the Fisher Transformation. This measure is applicable to neurons for which the CC maintained its sign between blocks. A positive ΔCC_indx_ indicates that the CC became stronger (larger in absolute value), whereas a negative ΔCC_indx_ indicates that the CC became weaker (smaller in absolute value). Plotting the histogram of ΔCC_indx_ between all pairs of non-target neurons that maintained their sign (n=1413, Figure 3D, bottom) showed that the large-scale changes in the pairwise correlations were relatively balanced between increases and decreases in amplitude, with a slight skew toward decreases (the median ΔCC_indx_ was −0.03). This result differs from the CC amplitude changes with the target neuron, which were strongly skewed toward an increase in CC amplitude (Figure 3D, top).

Overall, these results reveal a conservation (or balance) principle during BMI learning: while the increase in firing rate of the target neuron resulted in increased amplitude of CCs for pairs involving this neuron, overall compensation occurred in both firing rates and CCs across the population, such that changes in both these measures remained relatively balanced during BMI learning.

### A network model for exploration and learning

We next explored the computational principles underlying the changes in network activity during learning. A successful computational model of the learning process should account for all (or most) of the observed changes in neural activity patterns due to learning, as detailed above. In particular, the model should exhibit the capacity for exploration, a process that is essential for learning and requires strong variability in network activity. In cortex, such variability can emerge in networks operating in the regime of balanced excitation and inhibition [25,26,28,31]. Additionally, in such networks, the dynamics are expected to compensate for changes in firing rates and correlations, which could account for the relatively symmetric changes in firing rates (Figure 3A) and CC amplitude (Figure 3D, bottom) observed in the experiment. Because learning may introduce stability perturbations that can drive an EI network out of the balanced regime, we developed a novel model consisting of strong fixed recurrent connectivity, which maintains the balanced state, and weak plastic feedforward connectivity, which enables learning (Figure S3). The feedforward synapses learn using a simple, physiologically plausible, reward-dependent plasticity rule:

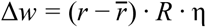

where Δ*w* is the change in strength of all the neuron’s feedforward input synapses,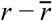 is the mean-subtracted firing rate of the target neuron, *R* characterizes the presence of reward, and η is a scalar scaling factor (the learning rate). Such a rule, termed the activity-reward covariance rule, has been proposed to explain decision-making processes [32].

We conducted simulations using the model (further described in Methods) to test whether this architecture and learning rule could account for all the observed changes in the experimental data. In a typical simulation run (Figure 3E-H), the target neuron increased its firing rate after the learning procedure began (Figure 3G). Firing rate changes across the population were relatively symmetric between increases and decreases (Figure 3E). Changes in firing rate (ΔFR_indx_) strongly depended on the pre-learning correlation patterns with the target neuron: negatively correlated neurons decreased their firing rate on average, while positively correlated neurons increased their firing rate on average (Figure 3F). The difference between these two groups was significant (p<10^-60^, Wilcoxon rank-sum test). The average firing rate of neurons that had no significant correlation with the target neuron did not change significantly (p>0.21, Wilcoxon signed rank test).

We also examined the correlations between non-target neurons and the target neuron in the simulation. The changes in these CCs between the observation-state and BMI-state blocks were similar to the changes found in the experimental data (Figure 3H). The correlation between these CCs during the observation- and BMI-states was 0.7 in the simulation (0.62 in data) and the percentage of CCs that did not change their sign in the two periods was 77% (72% in data). These CCs were mostly strengthened, as evidenced by the right-skewed distribution of ΔCC_indx_ (Figure 3H, top; p<10^-90^, Wilcoxon signed rank test), unlike the distribution of ΔCC_indx_ between non-target neurons, which was more symmetric and skewed to the left (Figure 3H, bottom). Further analysis exposed additional similarities between the data and the model (Figure S4). Finally, we tested the robustness of the model by running the simulation 100 times with different random seeds, which determined the recurrent network connectivity, the initial feed-forward weights, the initial network activity, the choice of target neuron and the arrival times of the feed-forward signal. The model exhibited highly robust behavior on all reported measures (Figure S5).

## Discussion

This study combines experiments and modeling to investigate the nature of brain-machine interface learning, in which subjects were required to volitionally increase firing rate of a single neuron in motor cortex to achieve reward. We report four major findings. (1) The neural correlation structure in a local circuit in motor cortex changes significantly depending on the action-perception state (Fig 2A,B). (2) The neural state during learning (BMI-state) was strongly related to the neural state preceding learning (observation-state), rather than to the neural state during arm movements (movement-state). Further, the correlation structure during the observation-state predicted learning-related changes in firing rates and pairwise correlations during the BMI-state. (3) Learning to increase the firing rate of a single neuron resulted in large-scale, symmetric changes in activity, which were not directly required by the BMI behavioral task. (4) A reinforcement learning model of a chaotic network operating in the regime of balanced excitation and inhibition predicted these experimentally measured learning-related changes.

Taken together, the experimental and modeling results support the idea that ongoing activity in the cortex represents internal brain states that evolved during different contexts, experiences, and sensory information, leading to selection of actions based on the current state and prediction of the consequences of the selected action. The model suggests a possible computational mechanism of learning, in which the input synapses to the local circuit are finely tuned to continuously adjust the relationships between the current inputs and the output, reflecting the optimal action-perception relationship at a given time.

### Relationship to reinforcement learning theory

We also propose a mechanistic model of how reinforcement learning is implemented in local cortical networks (Figure 4). The chaotic behavior of the network produces variability, which supports exploration. Concurrently, the AP state affects the correlation structure of the population. Thus, the exploration is probabilistically constrained to a subspace of neural activity state space (e.g., regions where strongly correlated neurons are active concurrently). Reward-modulated plasticity then guides this exploration by reinforcing activity patterns that lead to reward, leading to learning of the new AP state. This model explains the relationship between the pre-learning correlation patterns and the learning-related changes in firing rates. During exploration, activity patterns that increase the activity of the target neuron lead to reward, reinforcing this increase in activity. Then, neurons with positive correlations with the target neuron also tend to increase their firing rate, while negatively correlated neurons tend to reduce their activity. Because these changes lead to reward, they are also reinforced. In this way, we posit that the AP state affects the learning process via the neuronal correlation structure it establishes. This interpretation is consistent with findings in mice that strong spontaneous positive correlations with the target neurons were related to changes in firing rates [33].

**Figure 4.**
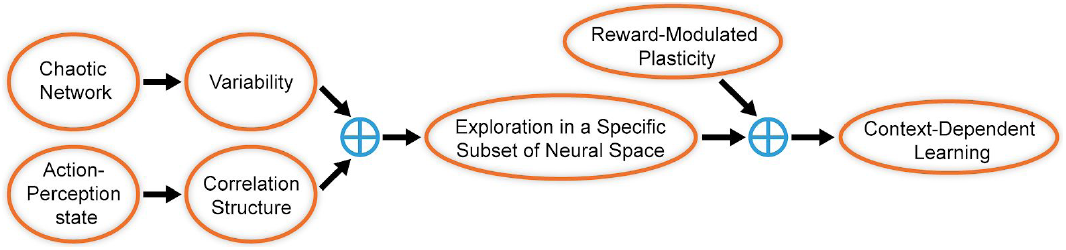
Flowchart depicting the processes that lead to context-dependent learning in our model. Variability is achieved through the chaotic dynamics of the network, while the action-perception state determines the functional correlation structure in which the network operates. This results in exploration that is limited to specific regions of neural space. When learning starts, reward-modulated plasticity reinforces activity patterns within that subspace that lead to reward, resulting in a learning process modulated by the action-perception state.

Previous computational models of volitional modulation of single-unit activity [34,35] used Hebbian learning to change the activity of the target neuron. However, they required the injection of external noise into the network to obtain variability. In contrast, variability is an inherent property of our model [26,28]. Further, our modeling results extend previous studies to account for the statistics of firing-rate changes of the non-target neurons, changes in correlations between the target and the non-target population as well as between non-target neurons themselves, and the dependence of firing rate changes on the pre-learning state. In our simulations, none of these features were forced to correspond to their homologues in the experimental data. Rather, they emerged as a natural consequence of a network operating in the balanced regime. Thus, our model captures key elements of the network dynamics as experimentally demonstrated, significantly beyond those features specifically required by the learning task.

### Relationship to previous studies of volitional neural control

A previous study [18] found that learning is easier when the task does not require systematic changes in the pre-learning correlation structure. Our RL framework accounts directly for those results. Our model and data show that the initial network exploration is indeed constrained by the pre-learning neuronal correlation structure. However, the authors suggested that the correlation pattern is fixed and therefore imposes a fixed constraint on learning. Our results and model show that the neural correlation structure that constrains the learning process is not fixed; instead, it can change dramatically depending on the AP state that precedes learning (Fig 2A). In addition, we find that the correlation structure indeed represents the initial condition of learning, but it adapts quite rapidly (few minutes) during learning to meet task requirements (Fig 3D).

### Significance for computational theories of cortical activity

Balanced networks were first suggested as a theoretical explanation for the irregular firing of cortical neurons [25,26] and later shown to account for other experimental observations, such as low correlations in cortical networks [31] and the emergence of orientation selectivity in V1 [36]. Our model shows how the balance of excitation and inhibition provides an intrinsic source of variability, resulting in explorative neuronal dynamics, which can drive a local reward-based learning rule to accommodate a variety of features of learning in local cortical networks. More specifically, we showed that learning in a balanced network results in relatively symmetric increases and decreases in firing rates across the population, which can explain the observed changes in the pairwise correlation pattern. The observed symmetric changes during learning highlight a homeostatic mechanism that sustains the balance of excitation and inhibition and thus maintains relatively stable overall network activity. To the best of our knowledge, this is the first study to incorporate behavioral learning in chaotic balanced network models.

### Implications for the clinical use of brain-machine interfaces

The development of clinical applications of invasive and non-invasive BMIs has followed two main approaches. The first aims to use brain activity to control external devices. These interfaces usually apply algorithms that interpret the neuronal activity to serve as the interface between the brain and the external device [3,4,6–8]. The second approach, sometimes termed operant neural conditioning, uses algorithms that are designed to induce changes in activity, and aim to assist the patient in generating specific neural activation patterns that can ameliorate cognitive deficits or engage plasticity mechanisms in the brain to restore appropriate neural connectivity [21,37,38]. Subjects can control a wide variety of neural patterns [11,15], but it remains unclear how to elicit specific activation of neurons not *directly* conditioned by the task, which limits the potential clinical use of this technique. Here, we were able to predict the effect of conditioning on the *non-target* neurons, suggesting that recording of the pre-conditioning state is beneficial for the design of the specific learning task (e.g. conditioning of single units with suitable correlation patterns), in order to elicit the desired activity patterns *throughout* the network during volitional control.

## Conclusion

In the present work, we operantly conditioned the activity of a single neuron and let the network learn with no additional constraints. At the core of the learning process, we found widespread reorganization across a population of neurons, including modifications of their firing rates and correlation patterns, which led to successful performance (increased firing rate of the target neuron and reward by the BMI). The model we used provides a possible explanation for the computational significance of the observed neuronal changes, and presents a framework that can encompass a wide variety of learning paradigms. The evolution of specific neural outcomes supported by general adaptation in the local network provides a powerful and robust substrate for learning in local cortical circuits. This attribute may be harnessed in future BMI settings to reshape the local network in a specific manner, for both basic science and clinical applications.

## Acknowledgments

We thank Y. Burak and H. Ito for helpful comments, A. Raz for help with the surgeries, and S. Freeman and A. Shapochnikov for technical assistance. This work was supported in part by the Bi-national Science Foundation (BSF), the Rosetrees Trust, the Gatsby Charitable Foundation, the Ida Baruch fund, and the Jack H. Skirball Fund in Brain Research. B. Engelhard was partly supported by the Felix M. Katar fund and the Khazzam-Goren fund. This work is dedicated to the memory of Dr. Dmitry Davidov.

## Methods

### Animals and Surgical Procedure

Two monkeys (Macaca fascicularis) were chronically implanted with a microelectrode array (Blackrock Microsystems) in the arm region of M1 contralateral to the performing arm, under anesthesia and aseptic conditions. Animal care and surgical procedures complied with the National Institutes of Health Guide for the Care and Use of Laboratory Animals and with guidelines defined by the Institutional Committee for Animal Care and Use at the Hebrew University.

### Behavioral Task

A single recording session was composed of four blocks. The first and last (movement-state blocks) consisted of a center-out task, with grip-and-reach movements to eight targets located in the corners of a three-dimensional cube. The monkeys used a robotic arm (Phantom Premium 1.5 High Force; SensAble Devices) and a custom-made gripping handle to control the movements. Grip force and the manipulandum position were sampled at 100 Hz. The monkeys could not see their arms. Instead, images of the targets and cursor were projected to the arm workspace. The cursor represented the hand location and grip force. Targets were defined as spheres of 8 mm radii, and the distance between the center of the origin and the center of each target was 4.85 cm. The trial began with a presentation of the origin in the center of the workspace. The monkeys were required to move the cursor to the origin, press the handle to a minimum force level and maintain the cursor position for a random period between 800 and 1300 ms (hold period). After the hold period, one of 8 targets selected randomly was presented, and the cursor had to remain in the origin for a second random period between 800 and 1300 ms (target presentation period). The origin then disappeared, which represented the go signal for movement. The monkeys then had to move the cursor to the target while maintaining the minimum grip force level (movement period). After reaching the target, the cursor needed to remain on the target for an additional 800 ms, after which a liquid reward was delivered and the trial ended. After an inter-trial interval of 1500 ms, a new trial began.

The second block of the session was termed the observation-state block (Figure 1A). In this block, the cursor and target were displayed in similar sizes and distances as in the movement blocks, but the monkeys’ arms were restrained, and the cursor position was determined randomly every 100 ms, resulting in random delivery of the reward. Four small red spheres located in the corners of the workspace were lit throughout this block to indicate the observation-state. In the BMI-state block (third block of the session), conditions were the same as in the observation-state, but the cursor position was determined by the mean firing rate of a single neuron (termed the target neuron). The details of the conditioning algorithm are described below. In the both the observation- and BMI-state blocks, an interpolation method was used to provide the appearance of smooth cursor movement.

### Electrophysiology

The recording array contained 100 electrodes (Blackrock Microsystems), of which 96 were functional, arranged in a 10×10 matrix with a 400 μm interelectrode distance. Spikes were extracted from the raw signal, sampled at 30 KHz, manually sorted using the histogram peak count algorithm and collected using the Cerebus data acquisition system (Blackrock Microsystems). A total of 34 recording sessions were analyzed (23 from monkey M, 11 from monkey Q). Of these, 24 sessions (16 from monkey M, 8 from monkey Q) were considered successful (see below) and were analyzed in further detail. A custom accelerometer (based on the MXR9500G/M chip, MEMSIC, Inc.) was placed on the middle finger of the contralateral hand of both monkeys during the observation- and BMI-state blocks and sampled at 100 Hz. Jerk amplitude was obtained by subtracting the mean from each of the accelerometer channels and calculating the square root of the sum of the squared first-order derivatives of each channel.

### Conditioning Algorithm

In each session, a single unit (from an electrode not previously chosen) was randomly selected for conditioning at the start of the BMI-state block. Firing rates were computed every 100 ms by counting spikes in a window of the last 400 ms. The cursor was positioned on the line between the origin and the target, with a linear correspondence to the firing rate. At 0 spikes, the cursor was placed at the origin. Above a maximum value the cursor was placed on the target, a reward was delivered, and the trial ended. The threshold for obtaining a reward was adjusted manually during the block to provide a relatively constant rate of reward, thus encouraging the monkeys to continuously improve their performance. A BMI session was considered successful if the ΔFR_indx_ (see below) was at least 0.4; however, this value usually was significantly higher (Figure 3A).

### State-space analysis for visualization of neural data

Gaussian Process Factor Analysis was implemented using a publicly available Matlab toolbox [29]. The procedure was conducted on the spike trains of the non-target neurons during successful trials of all blocks using a bin width of 50 ms. The resulting three-dimensional orthonormalized vectors for each trial were averaged to a single point, and plotted on a common scale for all blocks (Figure 2A). For clarity, every fifth trial is shown. This procedure was only used for visualization; all subsequent analyses were done on the full dimensionality of the data.

### Test for statistical significance of the difference in firing rates across blocks

Firing rates of the non-target neurons were estimated by summing spikes in a 1500-ms segment ending at reward for all successful trials in all blocks. In all sessions, we compared every pair of blocks in the following way: the distribution of neural distances within each block was estimated by calculating all the pairwise Euclidean distances between the different trials. Next, the distribution of neural distances between the blocks was estimated by calculating all the pairwise neural distances between trials belonging to different blocks. Finally, a Kolmogorov-Smirnoff test was run between the between-block distance distribution and a distribution composed of the union of the two within-block distances distributions.

### Test for statistical significance of the difference in neuronal correlations across blocks

To obtain the correlation matrices, we first convolved the spike trains of the non-target neurons with a zero-phase Gaussian kernel with s.d.=70 ms. Correlations between pairs of neurons were calculated for a concatenation of 5-second segments ending at reward for all successful trials in each block. Correlations were unrolled in a vector, and for each pair of blocks in a session the “between-blocks” correlation distance distribution was estimated by calculating the term-by-term root of the squared difference between the correlation vectors of the two blocks. Next, the “within-block” correlation distance distribution for each block was calculated as above but by comparing correlations found by using two disjoint and randomly sampled halves of the total number of trials. Finally, a Kolmogorov-Smirnoff test was run between the between-block distance distribution and a distribution composed of the union of the two within-block distance distributions.

### Quantifying the learning-related changes in firing rates and correlations

The change in firing rate index (ΔFR_indx_) between the observation-state block and the BMI-state block was calculated by first obtaining the mean firing rate in the 800-ms segment before reward in each block, and then dividing the difference between the two rates by their sum (see main text). For the ΔFR_indx_ involving movement blocks (Figure 3; Figure S1), firing rates for the movement block were taken from an 800-ms segment starting 200 ms before movement onset. Similar results were obtained if an 800-ms segment just before reward was used (not shown). To calculate the CCs of firing rates (Figures 3, S4, and S5), we binned the spike trains using 100-ms bins sampled every 50 ms, and calculated the cross-correlations of the resulting spike-count vectors across the whole block, allowing a maximum lag of 5 bins (250 ms). The CC was chosen as the correlation with lag that had the highest absolute value. The change in CC index (ΔCC_indx_) was calculated by subtracting the CC during the observation-state block from the CC during the BMI-state block and dividing the result by the sum of the CCs. This index was only calculated for CCs that maintained their sign in both blocks.

### Statistical test to assess whether firing rates during the BMI-state were more similar to those occurring during the observation-state or movement-state blocks

Mean firing rates were calculated in different segments of 800 ms along the trial for the different blocks. In the BMI-state block, firing rates were calculated just before reward. In the movement-state blocks, firing rates were calculated in 4 segments: just before reward, starting at the initial hold period, starting at the time of target presentation, and starting 200 ms before movement onset. These segments were chosen because a different pattern of neural activity was expected in each [39]. For the observation-state block, two segments were used: just before reward and at the start of the trial. To calculate neural distances from the non-BMI blocks to the BMI-state block, we first obtained activity vectors for all non-target neurons for each segment, using trials to each target direction separately. For each activity vector (for each segment and cursor direction), we calculated the normalized Euclidean distance from the activity vector of the BMI-state block using the square root of the sum of square differences, divided by the number of neurons. In order to account for possible general fluctuations in the mean activity, an optimal gain factor was found and applied to each activity vector, so as to minimize the normalized Euclidean distance by using the least-squares criterion. From the obtained distances, we selected the vector belonging to the target direction with the shortest distance for each segment. Thus, this analysis took into account the possibility that the monkeys used a movement-based re-aiming strategy during the BMI-state [40]. To account for the different number of trials in the different conditions (block and target direction), we found the condition with the lowest number of of trials (tmin) and then for all conditions randomly sampled tmin trials (from all trials in that block and with that target direction) to compute the firing rates. This analysis was repeated for 1000 iterations where, in each, the subset of tmin trials used for each condition was re-selected randomly.. For each iteration, we calculated a binary variable that was assigned a value of 1 if the distances were shorter than the “most-similar” (shortest distance) observation-state segment or 0 if the shortest segment was the most similar movement-state segment (from either of the movement-state blocks). The similarity index (Figure 2C) was the average of this binary variable. The chance level was 0.2 because we tested 8 segments from both movement-state blocks and 2 segments from the observation-state block.

### Statistical test to assess whether correlations during the BMI-state were more similar to those occurring during the observation- or movement-state blocks

To obtain the correlation matrices, we first convolved the spike trains of the non-target neurons with a zero-phase Gaussian kernel with s.d.=70 ms. Correlations were calculated in 5-second segments either ending at reward (all blocks) or beginning at the start of the trial (observation- and movement-state blocks) for trials to each target direction separately. Correlations were unrolled in a vector, and the sum of the squared differences was calculated between correlations in the BMI-state blocks and all the segments in all the other blocks. To account for the different number of trials in the different conditions (block and target direction), we found the condition with the lowest number of of trials (tmin) and then for all conditions randomly sampled tmin trials (from all trials in that block and with that target direction) to compute the correlation matrices. This analysis was repeated for 1000 iterations where, in each, the subset of tmin trials used for each condition was re-selected randomly. For each iteration, we calculated a binary variable that was assigned a value of 1 if the difference between the correlations vector of the BMI- and observation-state (in the observation-state condition where this difference was smallest) was smaller than the difference between the correlation vector of the BMI- and movement-state (in the movement-state condition where this difference was smallest, from either of the movement-state blocks). The similarity index (Figure 2D) was calculated as above. Here the chance level was 0.33 because we tested 4 segments from both movement-state blocks and 2 segments from the observation-state block.

### Analysis of variance

Analysis of variance was carried out using published matlab code [30]. In each trial, average activity was computed using 1 s of data (starting 250 ms before movement onset in the movement-state blocks, and 1 s before reward delivery in the observation- and BNI-state blocks; similar results were obtained if we used 1 s before reward in the movement-state blocks). We compared the matrices containing the average activities of all non-target neurons in the observation- and BMI-state blocks, or the first movement-state and the BMI-state blocks. In each session, we used the minimum number of dimensions that accounted for 95% of the variance of the data (concatenating all the blocks), according to a principal component analysis, and used 10000 random seeds to calculate the alignment to random spaces.

## Network model simulations

### Architecture and dynamics of the recurrent balanced network

We simulated a recurrent network composed of *N*_*E*_ excitatory (*E*) and *N*_*I*_ inhibitory (*I*) neurons, where each neuron receives inputs from an average of *K*_*E*_ excitatory and *K*_*I*_ inhibitory neurons. The input field of the *i*’th neuron in population α (α, β ∈ {*E, I*}) obeys the following dynamics:

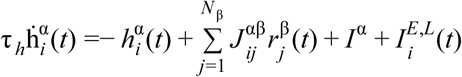

where τ_*h*_ is a time constant,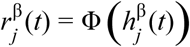 where Φ (*x*) is a non-linear transfer function, is the strength of the connectivity between the pre-synaptic *j*’th neuron in population β and the post-synaptic *i*’th neuron in population α, and *I*^α^ is a constant input drive to population α. We followed the same architecture as the well-studied balanced networks [25,26], such that 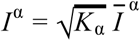 and the recurrent connections are modeled by a sparse and random interaction matrix; i.e.,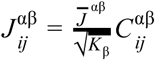 where 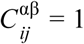 with probability 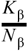 and 0 otherwise.

The behavior of this rate model (without the additional 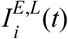 input) has been extensively explored [28] and was found to be equivalent to a full spiking neuronal model when the time constant τ_*h*_ (interpreted as a synaptic time constant) is not too small (see Model Parameters below); in our model, we add an additional feedforward (FF) input only to the *excitatory* neurons in the recurrent network which represents the command signal sent to motor cortex: 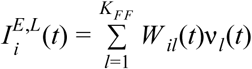, where v_*l*_ (*t*) is the firing rate of the *l*’th neuron in an external network and *K*_*FF*_ (assumed to be large) is the number of FF connections each neuron in the recurrent network receives from this additional FF input. The strength of these FF connections alone changes throughout learning and scales with the number of connections; i.e.,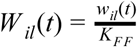 where *w*_*il*_ are order unity. These FF synapses are weak compared to the recurrent synapses because the sum of FF synapses for a given neuron is order unity, whereas the sum of recurrent synapses is order 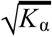. This implies that the command signal will not change the global state of the network; i.e., the mean of the firing rates and correlations. In particular, it guarantees that the network remains throughout learning in the balanced regime. In this work, we used Φ (*x*) as Φ (*x*) = {0 | *x* ≤ 0; *x* | 0 < *x* < *d*; *S*(*x*) | *x* ≥ *d*}, where 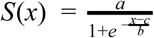. Similar results were obtained using the threshold-linear function: Φ (*x*) = *max*(0, *x*) (not shown).

### Learning procedure

We use a covariance-based learning algorithm to increase the firing rate of the target neuron. The rationale behind this family of learning rules is that synaptic weights are changed according to the co-variation of a global reward signal with local neuronal activity, such that a small increase (decrease) in the expected reward that is correlated with an increase in the neuronal activity will be implemented (discarded) in the system. Specifically, we used the following learning rule:

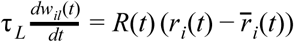

where 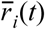 is an exponential running average with a time constant of 10 s, and 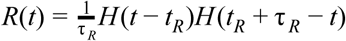 is the delivery of the reward following time *t*_*R*_ for a duration of τ_*R*_, where *H*(*x*) is the Heaviside step function. We assume that the FF input comes from an asynchronous network with a population average firing rate of 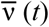, which changes according to the command signal sent to motor cortex (see below). As the activity of neurons in the FF network is uncorrelated with the strength of their projections, the FF input is simplified to 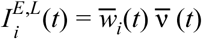, where 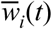 is the expected value (over synapses) of the FF synapses for the *i*’th neuron. Thus, the learning rule is reduced to the dynamics of the mean connectivity strengths: 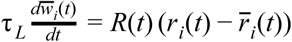.

The initial weights 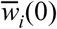 were randomly sampled from a standard normal distribution. We took an instantaneous delivery of reward, such that τ_*R*_ =1 ms. In the simulation, at the beginning of the BMI block, one excitatory (target) neuron was chosen randomly for conditioning (as in the experiment); a reward was delivered whenever the rate of the target neuron (smoothed with an exponential running average with a time constant of 400 ms) increased its firing rate with respect to an exponential running average with a time constant of 10 seconds. A reward was delivered no sooner than 1500 ms after the previous reward, to mimic the inter-trial interval in the experiment.

### Simulation of the incoming feed-forward (volitional) signal

We simulated the average firing rate of the feed-forward projections 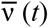 as an episodic signal manifesting at random times as a consequence of the command signal. Each episode of increased drive was modeled as a square wave with a duration of 300 ms and an amplitude of 2.5 that was smoothed by a zero phase Gaussian filter with s.d.=20. When a given episode ended, the start time of the next episode was sampled randomly from a uniform distribution between 500 and 2000 ms.

### Maintaining the balanced regime during learning

As stated above, because the synapses of the command feedforward projections are weak compared to the recurrent synapses, the balance between excitation and inhibition, which is a dynamic property of the recurrent network, is maintained for a wide range of network parameters [26,28]. In this sense, the network behavior is robust to the specific choice of parameters. These parameters do, however, determine the specific values of the population-average firing rate of the neurons, which can be calculated by solving a set of equations expressing the fact that the network operates in the balanced regime [26]. The solution to these equations does not depend on the volitional input, as the latter is much weaker than the background (constant drive) input or the recurrent synapses. In our model, we chose the network parameters to fit the average firing rates estimated from the experimental data (see below) and to position the network in the chaotic balanced state. As long as the network is in that state, the simulation results are robust to the specific set of parameters. In such conditions, increases in firing rates in one subpopulation induced by the learning process must be compensated across the network by a decrease in firing rates in a different subpopulation. The same argument is valid for the observed balance in the changes in neuronal correlations [31].

### Model parameters

The autocorrelations of the neurons in the model are mainly determined by τ _*h*_. We therefore set τ_*h*_ = 20 ms, which corresponds to a decay of ∼ 400 ms of the autocorrelation, as typically found in the data during the observation-state blocks (not shown). As for the network size and number of connections, we used the fact that the distribution of correlations in balanced networks has finite width when considering a finite number of connections and neurons in the network to ensure that correlations in our simulated network were similar to the estimated correlations from the data. We therefore set *N*_*E*_ = 4800, *N*_*I*_ = 1200 and *K*_*E*_ = *K*_*I*_ = 200. We chose the learning time scale to fit the typical time it took the monkeys to learn the task: τ_*L*_ = 630 s. The average firing rate of the neurons was fixed by using the balance equation [26] to fit the estimated firing rates from the data, by using the following parameters: 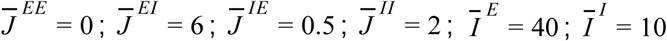. These parameters provide an average global firing rate close to 7 Hz, up to corrections resulting from using a finite number of connections [26]. The average firing rate also remained fixed during learning, as described above. For the threshold-linear-sigmoid transfer function, the following parameters were used: *a* = 200, *b* = 40, *c* = 100, *d* = 30. The time step of the simulation was Δ*t* = 1 ms.

### Data analysis of the simulations

The simulated data were analyzed similarly to the experimental data (see Results and Methods). As our recordings were biased toward the large pyramidal cells in layer 5 of motor cortex, we assumed that the majority of our recorded neurons were excitatory and therefore analyzed only the excitatory cells in the simulation to enable the comparison. Additionally, because we did not detect any neurons with average firing rates below 0.1 Hz in the experiment, we assume this was an additional bias in our recordings, and thus did not analyze neurons in the simulation that had an average firing rate of less than 0.1 Hz in the pre-learning period. Finally, due to computational limitations, cross-correlations were computed for a random subset of pairs of non-target neurons.

