## Supplementary Information for "Neuronal activity and learning in local cortical networks are modulated by the action-perception state"

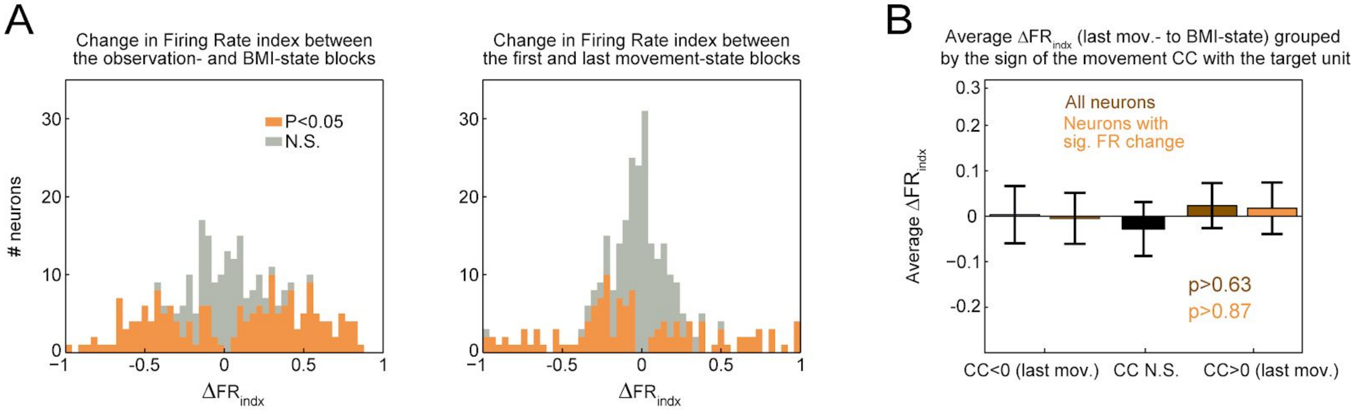


**Figure S1. Control Analyses**

(A) Histograms of $\Delta FR_{indx}$ shown in a common scale. Left: between the observation-state block and the BMI-state block (same as Figure 3A). Right: between the first and last movement-state blocks. The spread of the $\Delta FR_{indx}$ between observation- and BMI-state blocks (left) was significantly larger than that of the distribution of $\Delta FR_{indx}$ between the two movement-state blocks (right, p<2.3x10-5, Brown–Forsythe test).

(B) Average $\Delta FR_{indx}$ calculated between the last movement-state block (last mov.) and BMI-state block, grouped by the sign of the CC with the target neuron during the last movement-state block. The plot shows no relationship between the correlation pattern with the target neuron (during the last movement-state block) and the calculated $\Delta FR_{indx}$. Error bars are s.e.m.

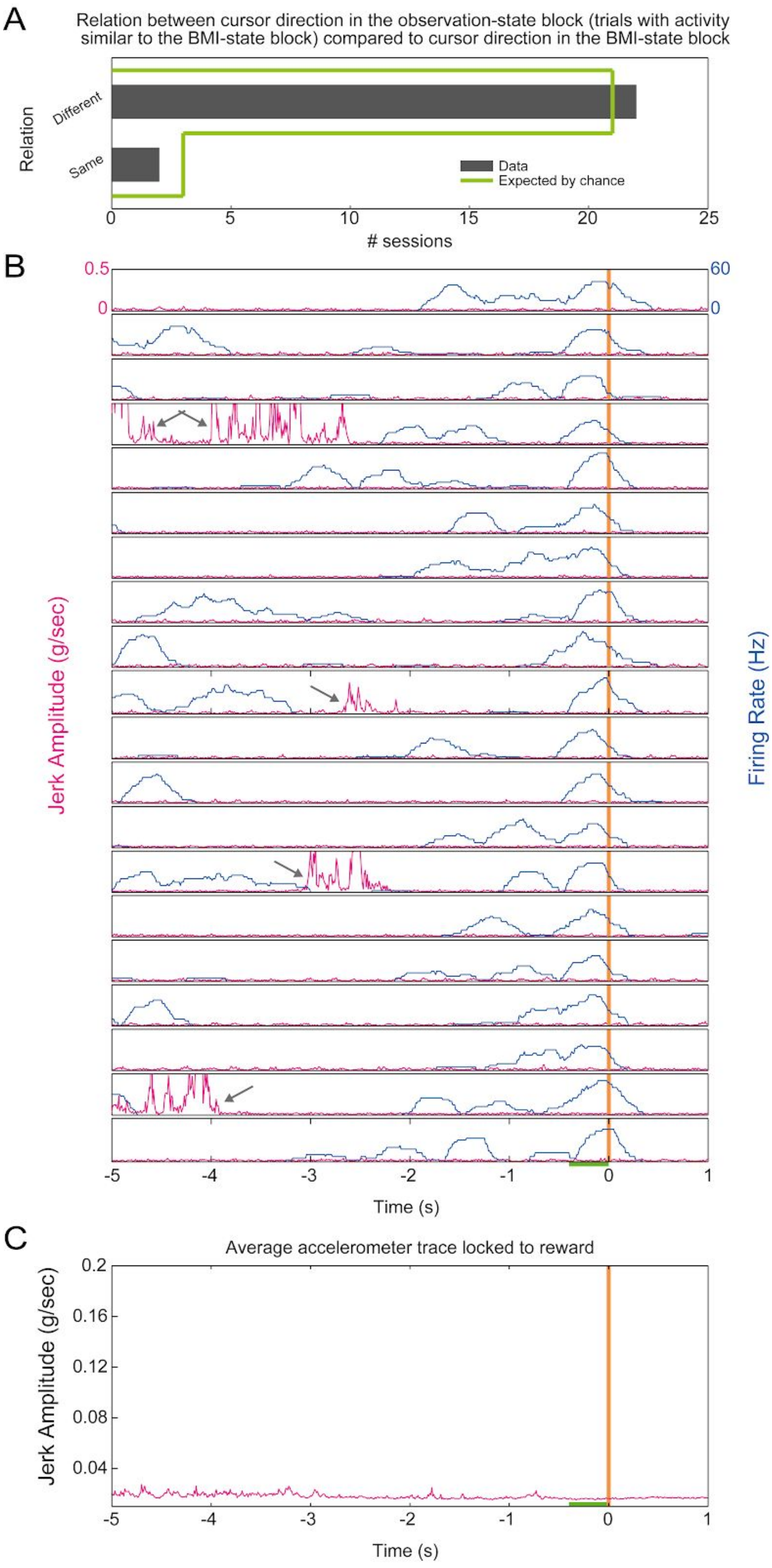


**Figure S2. Sensory- and motor-related responses do not account for the firing patterns during the BMI-state.**

(A) Relationship between the sensory stimulus present for similar neural patterns in the observation- and BMI-state blocks. Since activity in the motor cortex is modulated by sensory stimuli [1,2] we examined whether there was a strong sensory component to the activity patterns during the BMI-state block. During the observation-state block there were trials in which the cursor moved in 8 possible directions, whereas a single random direction was presented during the BMI-state. To test the possibility that the sensory stimulus had an effect on the observed neural patterns during the BMI-state, for each session we found the direction of the cursor in the observation-state for which the firing rates were most similar to those occurring during the BMI-state. If there were a strong sensory component to the BMI-state firing patterns, we would expect the "most similar" observation-state cursor direction to be the same as BMI-state cursor direction. However, we found no evidence for this. The graph shows the number of sessions where the "most similar" observation-state direction was the same (2/24) or was different (22/24) from the direction during the BMI-state. This ratio is not significantly different from chance ($p>0.76$, Binomial test). Thus, we conclude that the sensory component affecting the neural patterns during the BMI-state was small, if any.

(B) An accelerometer was placed on the middle finger of the contralateral hand of both monkeys during the BMI-state blocks (See Methods). The panel shows accelerometer traces during 20 consecutive trials of the BMI-state block from a single session. For each trial, data are shown from 5 s before to 1 s after reward delivery. Amplitude of the jerk is shown in magenta, and reward times are depicted by orange vertical lines. For reference, the firing rate of the target neuron (collected in 400 ms bins) is shown in blue. Both measures are shown in a common scale for all trials. The time window used to calculate the neural activity that led to reward is represented by the green line before time 0 (reward delivery). The gray arrows point to hand and finger movements that are clearly identifiable by the accelerometer traces, which did not lead to reward.

(C) Reward locked average accelerometer trace for the same session as in A. Reward was given at time 0, and the time window of the neural activity which led to reward is represented by the green line. The magnified scale (as compared to A) still shows no relationship between hand activity and reward. This analysis, coupled with the analysis of the video recordings of the head and body which did not show gross movements leading to reward strongly suggests that neural control of the BMI was achieved irrespective of physical movement, as has been shown in several previous studies in this type of task [3–6].

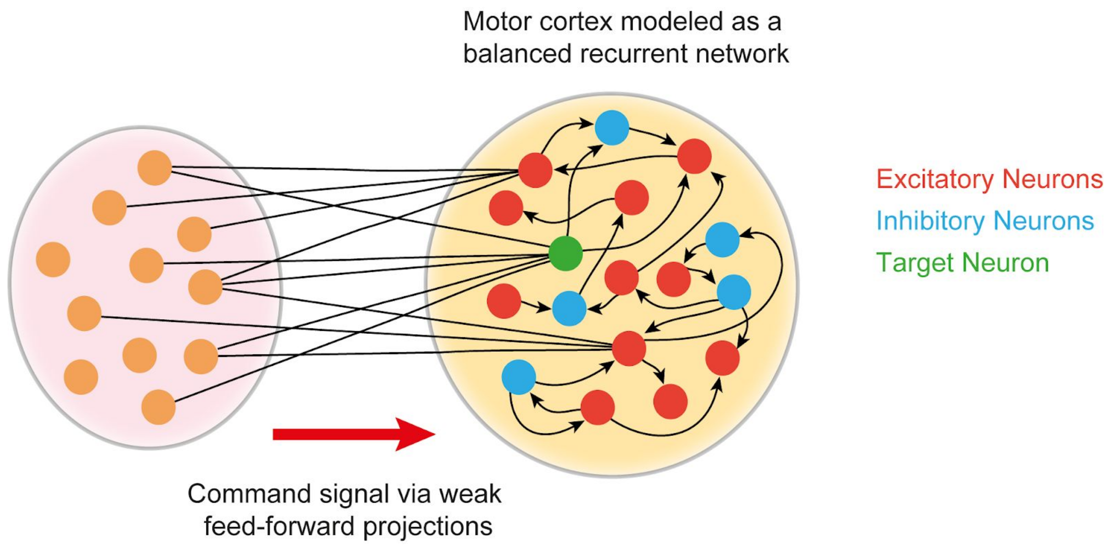


**Figure S3. Graphical depiction of the network model.**

The command signal reaches the motor cortex via weak feed-forward projections. The weights of these projections alone are modified by the activity-reward covariance rule described in the text. The motor cortex is modeled as a balanced recurrent network with 80% excitatory (red) and 20% inhibitory (blue) units. In each simulation run the target unit (green) was randomly selected from the excitatory population (see Methods for further details on the model).

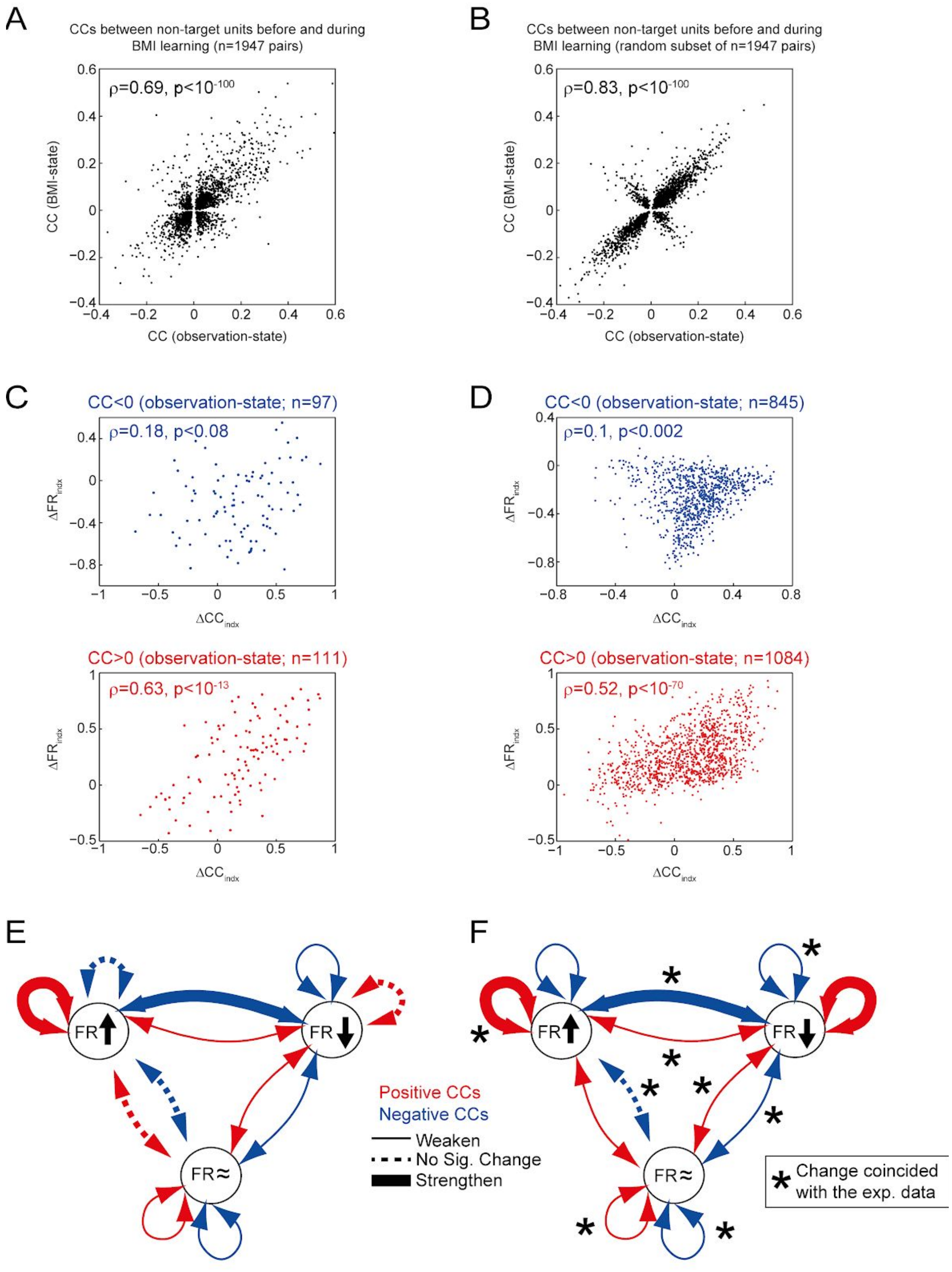


**Figure S4. Additional similarities between the data and the model.**

The figure shows several analyses performed on the data (left column) and the simulation results based on the model (right column; the same simulation run used in Figure 3 and the main text).

(A) Scatterplot of the CCs between pairs of non-target neurons in the observation- and BMI-states (experimental data).

(B) Scatterplot of the CCs between pairs of non-target neurons in the “observation”- and “BMI”-states (model simulation). The plot shows strong correlation between the two sets of CCS, similarly to that observed in the experimental data. To visualize the comparison with the experimental data in (A), a random subset of n=1947 was used for this plot (the overall correlation between the CCs in the two blocks for full dataset was also 0.83).

(C) Relationship between the change in CC amplitude with the target neuron ($\Delta CC_{indx}$)and change in average firing rate ($\Delta FR_{indx}$ ) between the observation- and BMI-state blocks (experimental data). This relationship was computed separately for neurons that had a negative CC with the target neuron (top; blue dots) or positive CC with the target neuron (bottom; red dots) during the observation-state block. These panels show that the relationship between the changes in correlation strength and firing rate depended on the sign of the CC with the target neuron: for negatively correlated neurons this relationship was weak, whereas it was strong for positively correlated neurons.

(D) Relationship between the change in CC amplitude with the target neuron ($\Delta CC_{indx)}$ and change in average firing rate ($\Delta FR_{indx}$ ) between the “observation”- and “BMI”-state blocks (simulation results). Similar to the experimental data, the relationship between the changes in correlation strength and firing rate depended on the sign of the CC with the target neuron: for negatively correlated neurons this relationship was weak, whereas it was strong for positively correlated neurons.

(E) In order to further investigate the changes in correlation strength between the pairs of non-target neurons in the experimental data, we first separated this population into three groups depending on whether the neurons significantly increased, decreased, or did not significantly change their firing rate between the observation- and BMI-state blocks. (denoted as FR↑, FR↓, and FR≈ respectively). We then quantified amplitude changes in positive and negative CCs within and between these groups by calculating the median $\Delta CC_{indx}$ for pairs of neurons in each subset (there were 12 subset of pairs given the division into positive or negative CCs and the 6 possible combinations within and between the 3 groups). A significant positive or negative median $\Delta CC_{indx}$ ($p<0.01$, Wilcoxon signed rank test) indicates that the CCs in the subset were strengthened or weakened respectively, whereas a non-significant median $\Delta CC_{indx}$ indicates no significant change in CC strength (see Figure S6 for the distribution of $\Delta CC_{indx}$ for all subsets of pairs). This panel illustrates all the interactions between the different groups and indicates the significant increases and decreases in amplitude: Positive (but not negative) CCs within the FR↑ group are strengthened, whereas negative (but not positive) CCs within the FR↓ group are weakened. Between these two groups, both modes take place: positive CCs are weakened and negative

CCs are strengthened. Thus there was an intricate adaptation pattern of the correlation structure between neurons that modulated their firing rate in relation to the task. Interestingly, we also observed changes in correlation strength between neurons that did not modulate their firing rates. Both positive and negative CCs within the FR≈ group were weakened. This highly statistically significant result ($p<7x10^{-4}$, see Figure S6) is by construction not driven by firing rate changes, and provides evidence for desynchronization between neurons that did not exhibit task-related activity.

(F) Similar to (E) but for the simulation results. The panel shows the changes in CC amplitude between all subsets of pairs (negative or positive CCs, within and between groups of neurons that increased, decreased or did not significantly changed their firing rate between the “observation” and “BMI” states, denoted as FR↑, FR↓, and FR≈ respectively). Out of the 12 subsets tested, the simulation matched 9 CC amplitude changes observed in the experimental data (denoted by asterisks in this panel. Assuming a uniform probability for the model and the data to match each CC amplitude change (which can strengthen, weaken, or not change significantly), matching of 9 out of 12 changes constitutes a p-value of 0.0033.

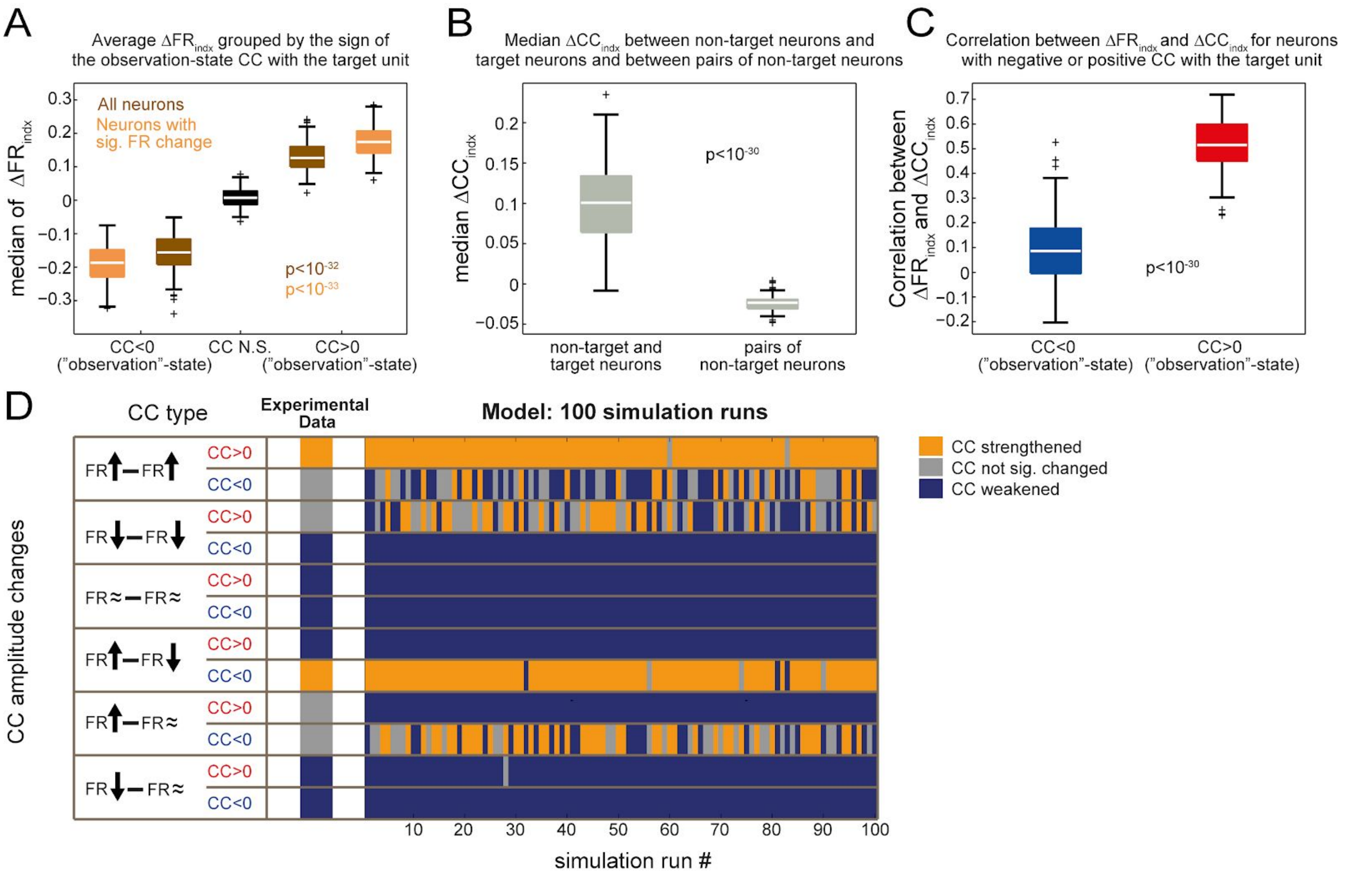


**Figure S5. Robustness of the balanced network model.**

To test the robustness of the model to a specific simulation instantiation (the recurrent network connectivity, the initial feed-forward weights, the initial network activity, the choice of target neuron and the arrival time of the feed-forward signal), we ran the simulation 100 times with different instantiations of these values. The results show robust similarities to the experimental data independent of the specific model instantiation.

(A) Boxplots of the average $\Delta FR_{indx}$ grouped by the sign of the "observation"-state CC for 100 simulation runs. Bottom and top edges of the filled boxes mark the 25th and 75th percentiles of the distribution; the median is denoted by a horizontal white line. Significance of the difference between the distributions (Wilcoxon rank-sum test, calculated separately for all units and for units which evidenced a significant change in their firing rates) is marked on the plot. Note the similarity to the sample simulation run shown in the main text (Figure 3E-H), and the experimental data (Figure 3A-D).

(B) Boxplots of the median $\Delta CC_{indx}$ between non-target and target neurons (left) and between pairs of non-target neurons (right) for 100 simulation runs. This panel summarizes the results shown in Figure 3H for all simulation runs.

(C) Boxplot of the correlation between $\Delta FR_{indx}$ and $\Delta CC_{indx}$ for non-target units with negative and positive CCs with the target neuron for 100 simulation runs. This panel summarizes the results shown in Figure S4D for all simulation runs.

(D) Summary of the CC amplitude changes found in 100 simulation runs. Each row denotes one of 12 subsets of CCs (legend on the left). In each run, CCs in each category can strengthen (orange) weaken (dark blue) or not change significantly (gray). For comparison, changes found in the experimental data are shown on the left. Note the consistent behavior of the model (matching the experimental data) for most interactions except for those where the data did not provide sufficient evidence of a significant change in CC amplitude (rows 2, 3 and 10).

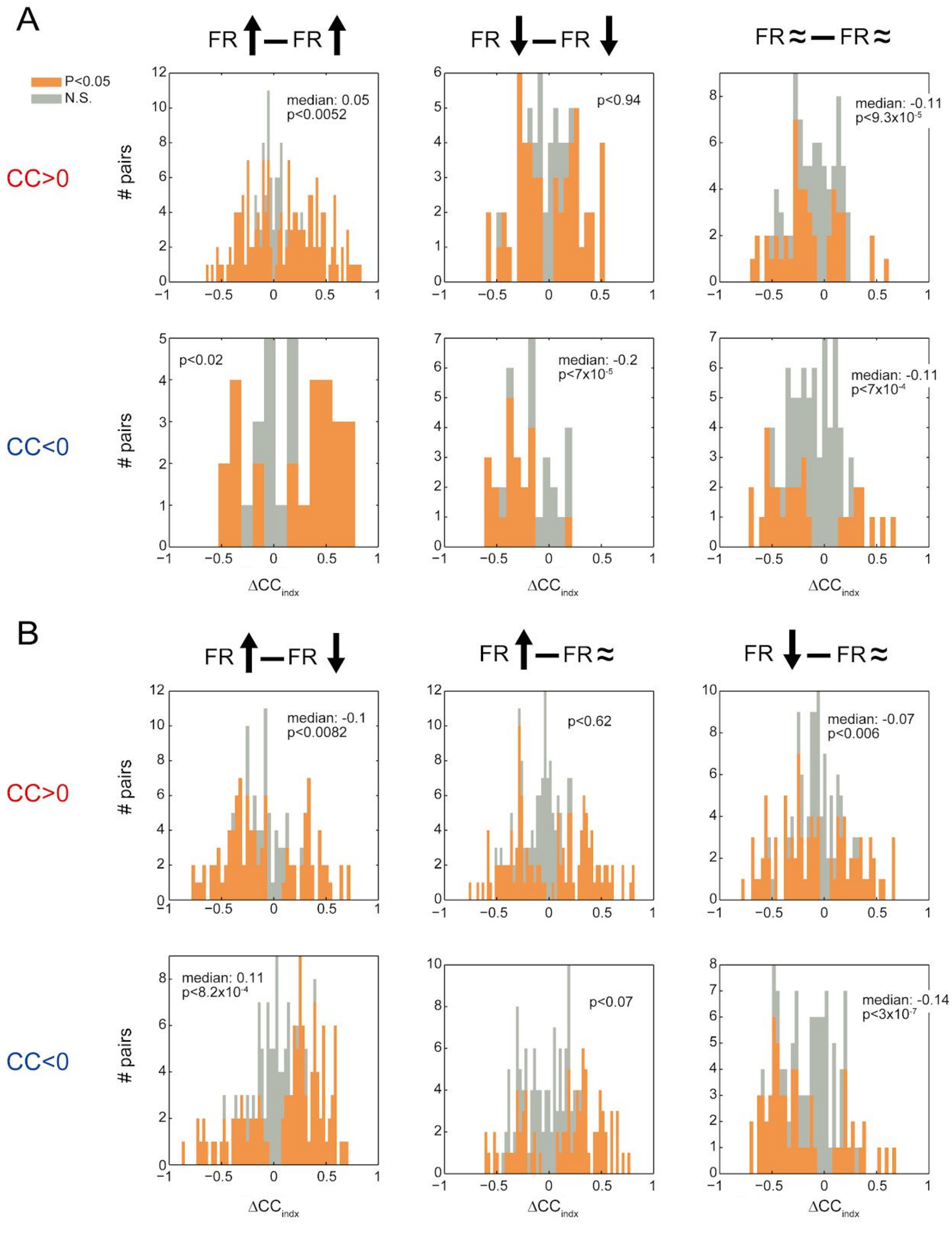


**Figure S6. Distributions of $\Delta CC_{indx}$ between pairs of non-target neurons.**

Each panel shows the distribution of $\Delta CC_{indx}$ between the observation- and BMI-state blocks for a subset of pairs. First, non-target neurons were divided into 3 groups: neurons that enhanced (FR↑) reduced (FR↓) or did not significantly change (FR≈) their firing rate between the observation- and BMI-state blocks. Next, these groups were used to test 12 subsets of pairwise CCs: positive or negative observation-state CCs between or within each of these 3 neuronal groups (see Figure S4 for a summary on this analysis).

(A) Histograms of $\Delta CC_{indx}$ for within group interactions, for positive (top) and negative (bottom) CCs (for all pairs that maintained the sign of their CC between the observation- and BMI-state blocks). Significance of the change in each CC ($p<0.05$, z-test on the Fisher transformed CCs) is color coded. In each panel the significance of the median is shown (Wilcoxon rank-sum test) as well as the median if significantly different from 0.

(B) Same as (A), for the between groups interactions.
